# Direct and Indirect Salt Effects on Homotypic Phase Separation

**DOI:** 10.1101/2024.05.26.596000

**Authors:** Matthew MacAinsh, Souvik Dey, Huan-Xiang Zhou

## Abstract

The low-complexity domain of hnRNPA1 (A1-LCD) phase separates in a salt-dependent manner. Unlike many intrinsically disordered proteins (IDPs) whose phase separation is suppressed by increasing salt concentrations, the phase separation of A1-LCD is promoted by > 100 mM NaCl. To investigate the atypical salt effect on A1-LCD phase separation, we carried out all-atom molecular dynamics simulations of systems comprising multiple A1-LCD chains at NaCl concentrations from 50 to 1000 mM NaCl. The ions occupy first-shell as well as more distant sites around the IDP chains, with Arg sidechains and backbone carbonyls the favored partners of Cl^−^ and Na^+^, respectively. They play two direct roles in driving A1-LCD condensation. The first is to neutralize the high net charge of the protein (+9) by an excess of bound Cl^−^ over Na^+^; the second is to bridge between A1-LCD chains, thereby fortifying the intermolecular interaction networks in the dense phase. At high concentrations, NaCl also indirectly strengthens π-π, cation-π, and amino-π interactions, by drawing water away from the interaction partners. Therefore, at low salt, A1-LCD is prevented from phase separation by net charge repulsion; at intermediate concentrations, NaCl neutralizes enough of the net charge while also bridging IDP chains to drive phase separation. This drive becomes even stronger at high salt due to strengthened π-type interactions. Based on this understanding, four classes of salt dependence of IDP phase separation can be predicted from amino-acid composition.

## Introduction

Biomolecular condensates formed via liquid-liquid phase separation (LLPS) mediate a variety of cellular functions such as biogenesis of the ribosome and stress response.^1,2^ The driver for phase separation or condensation is intermolecular interactions, including electrostatic, hydrogen bonding, π-π, cation-π, amino-π, and hydrophobic.^3–5^ Salt can tune all these interactions and thus exert significant effects on phase separation. While the screening effect of salt on electrostatic interactions is well-known, its effects on other types of interactions may be indirect and perhaps are less appreciated. In particular, high salt strengthens hydrophobic interactions by increasing the surface tension of water.^6,7^ Because salt can exert disparate effects on different types of interactions, it can be used as a perturbation to dissect the relative importance of these interactions in phase separation.^8,9^ Inside cells, proteins can encounter varying salt conditions at different locations or at different times, and therefore the drive for their phase separation can span a wide range.

Numerous studies of salt effects on protein phase separation have been reported. The most typical effect is the suppression of phase separation by screening electrostatic attraction, both for homotypic systems^10–18^ and for heterotypic systems.^8,19^ However, salt can also promote phase separation, although the mechanism is not always clear.^17,20–29^ In particular, Krainer et. al.^28^ observed a “reentrant” salt effect on the phase separation of five intrinsically disordered proteins (IDPs): phase separation occurs without salt, is prohibited by medium salt, and reemerges at high salt. They attributed the reemergence of phase separation to strengthened π-type and hydrophobic interactions that overcompensate weakened electrostatic attraction. A unifying understanding of how salt affects the phase separation of IDPs is still lacking. For example, it is an open question whether salt effects on phase separation can be predicted from the protein sequence. Deep knowledge, in particular at the atomic level, of how salt affects intermolecular interactions and, ultimately, phase separation is required.

The low-complexity domain of hnRNPA1 (A1-LCD) represents another IDP where an atypical salt effect was reported.^17^ The full-length protein, comprising folded domains along with the LCD, phase separates in the absence of salt and the tendency to phase separate is reduced upon adding salt, thus exhibiting the typical salt effect. A screening mechanism, specifically of electrostatic attraction between the LCD and the folded domains, is supported by small-angle X-ray scattering and coarse-grained molecular dynamics (MD) simulations. In contrast, A1-LCD does not phase separate without salt and starts to do so only after 100 mM NaCl is added. An earlier study revealed that π-types of interactions, mediated by aromatic residues, drive the phase separation of A1-LCD.^30^ A follow-up study, based on charge mutations, further showed that the net charge plays a strong suppressive role.^31^ The saturation concentration (*C*_sat_) for phase separation is minimum at a net charge near 0, and increases by two orders of magnitude, signifying an enormous weakening of the drive for phase separation, when the net charge moves away from neutrality in either direction. In a more recent study, salt promoted the homotypic phase separation of both A1-LCD and FUS-LCD, but suppressed the heterotypic phase separation of their mixture.^8^ The salt effect on the phase separation of A1-LCD was modeled by a Debye-Hückel potential in coarse-grained simulations.^32^ Similarly, the salt effect on the phase separation of another IDP, Ddx4, was analyzed using the random phase approximation based on a coarse-grained representation.^18,33,34^ Coarse-grained simulations with explicit water and ions have been used to study salt effects in both homotypic and heterotypic phase separation.^35^

All-atom MD simulations can uniquely provide mechanistic insight into the driving force and properties of biomolecular condensates.^5^ For example, these simulations showed that ATP, a small molecule with a -4 charge, bridges between positively charged IDP chains in driving phase separation.^36^ The intermolecular interactions quickly break and reform, explaining why the condensates can rapidly fuse despite very high macroscopic viscosity. Similarly, quick breakup and reformation of salt bridges in a heterotypic condensate allow the protein molecules to be extremely dynamic in a highly viscous environment.^19^ Recently, all-atom MD simulations provided explanations for wide variations in phase equilibrium and material properties among condensates of tetrapeptides with different amino-acid compositions.^37^ These and other simulations^38^ show that all attractive residue-residue contacts contribute to the drive for phase separation.

Here we study salt effects on A1-LCD condensation by all-atom MD simulations. The simulations reveal two direct effects and one indirect effect of NaCl: neutralization of net charge and bridging between protein chains as well as strengthening of π-type interactions by drawing water away from the interaction partners. We also present a unified picture of salt dependences of phase separation by defining four distinct classes and predict these classes from amino-acid composition.

## Results

### Salt condenses A1-LCD and increases inter-chain interactions

The 131-residue A1-LCD is comprised mostly of Gly and Ser (51 and 22, respectively), followed by 18 aromatic residues (11 Phe and 7 Tyr), 17 residues with sidechain amides (13 Asn and 4 Gln), and 15 charged residues (10 Arg, 2 Lys, and 3 Asp), with a large net charge of +9 (Figure 1A). Using initial conformations of A1-LCD from the previous single-copy simulations,^39^ we built an 8-copy model for the dense phase (with an initial concentration of 3.5 mM) at five NaCl concentrations ranging from 50 to 1000 mM (Figure 1B). This initial concentration is close to the measured concentration at the critical point,^30^ thereby facilitating the comparison between low salt (where phase separation does not occur) and high salt (where phase separation does occur). Each system was simulated in four replicates for 1.5 μs each. All the results reported here are averages over the four replicate simulations. From the same initial configuration with very few inter-chain contacts, the systems maintain the initial loose configuration at 50 mM NaCl (Figure 1C) but condense noticeably at 1000 mM NaCl (Figure 1D). At low salt, the protein chains have a tendency to fill the simulation box, occasionally spanning the entire box in one or two of the three orthogonal directions and exhibiting larger-scale reconfiguration based on root-mean-square-fluctuation (RMSF) calculation (Supplementary Figure 1). In contrast, at high salt, the chains aggregate into an ellipsoidal particle, with the RMSF reduced by 27%.

**Figure 1.**
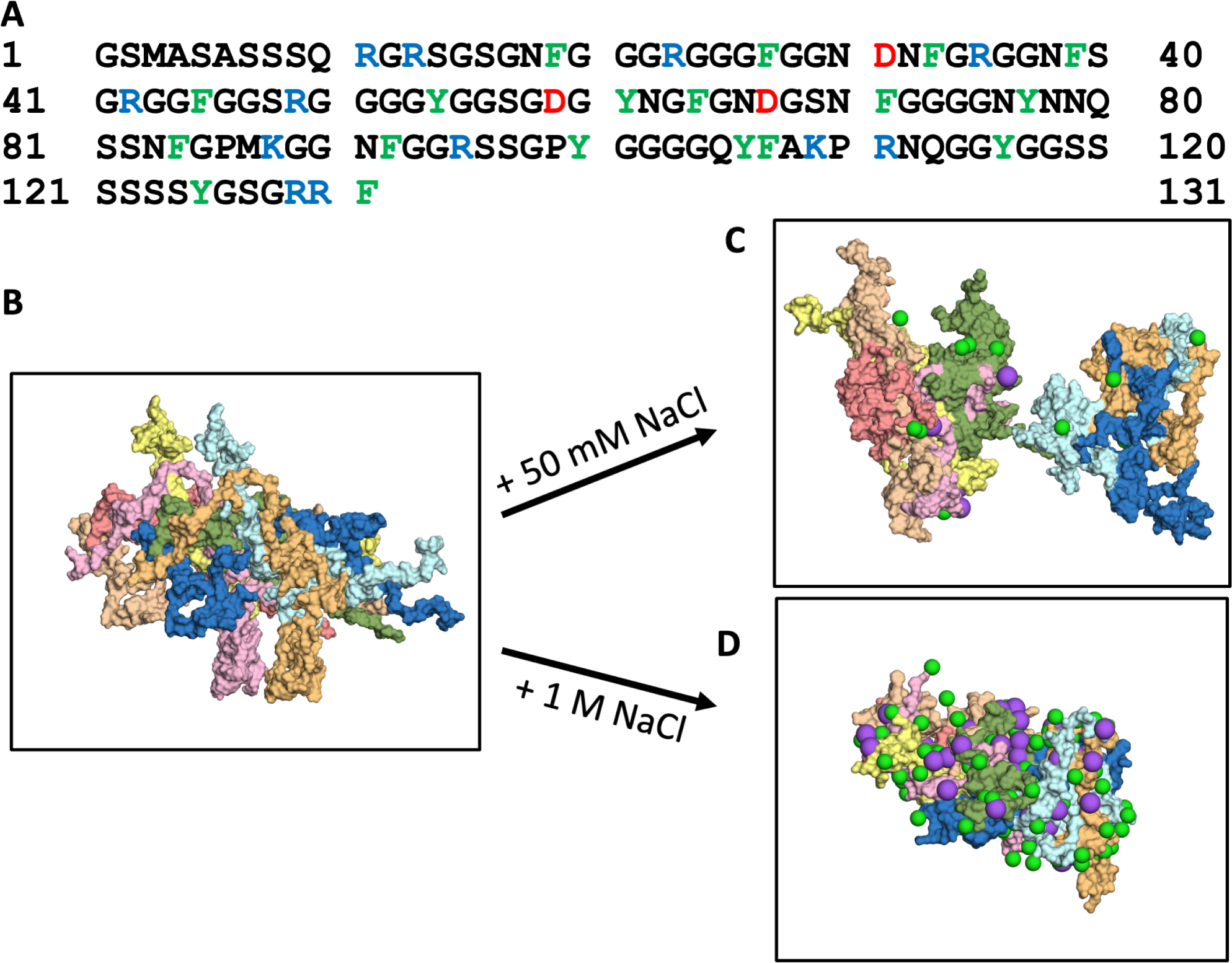
Amino-acid sequence of A1-LCD and molecular dynamics simulations of its condensation. (A) Amino-acid sequence. (B) First frame and (C, D) a frame at 1000 ns from 1.5-μs simulations of the 8-chain systems at low and high salt. In all figures, Cl^−^ or Na^+^ ions are represented by green and magenta spheres, respectively.

We also quantified the contrast between low and high salt by calculating *D*_max_, the maximum side length of the rectangular box that parallels the simulation box and circumscribes the multi-chain system. The *D*_max_ values are close to the initial value (∼150 Å) at low salt (50 mM NaCl), but decrease to ∼135 Å at intermediate salt (150 and 300 mM NaCl) and further to ∼120 Å at high salt (500 and 1000 mM NaCl; Figure 2A). Overall, the systems show successive condensation on going from low salt to intermediate salt to high salt. Modeling the condensed particle as an ellipsoid with the principal diameters given by the maximum dimensions in the three orthogonal directions, we can estimate its concentration to be 23 mM, which is similar to the measured concentration in the dense phase.^30^ A small part of the reason for the growing condensation is the compaction of the individual chains. The average radius of gyration (*R*_g_) shows a decreasing trend with increasing salt (Figure 2B), which matches the experimental data.^17^ Correspondingly, the average number of intrachain contacts per residue increases slightly, by 5%, when the salt concentration is increased from 50 mM to 1000 mM (Figure 2C). However, the main driver of the condensation is inter-chain interactions, with inter-chain contacts per residue increasing by 23% over the same salt range. Clearly, high salt induces A1-LCD condensation, with an increased number of interactions between protein chains. Below we present the molecular mechanisms for these effects.

**Figure 2.**
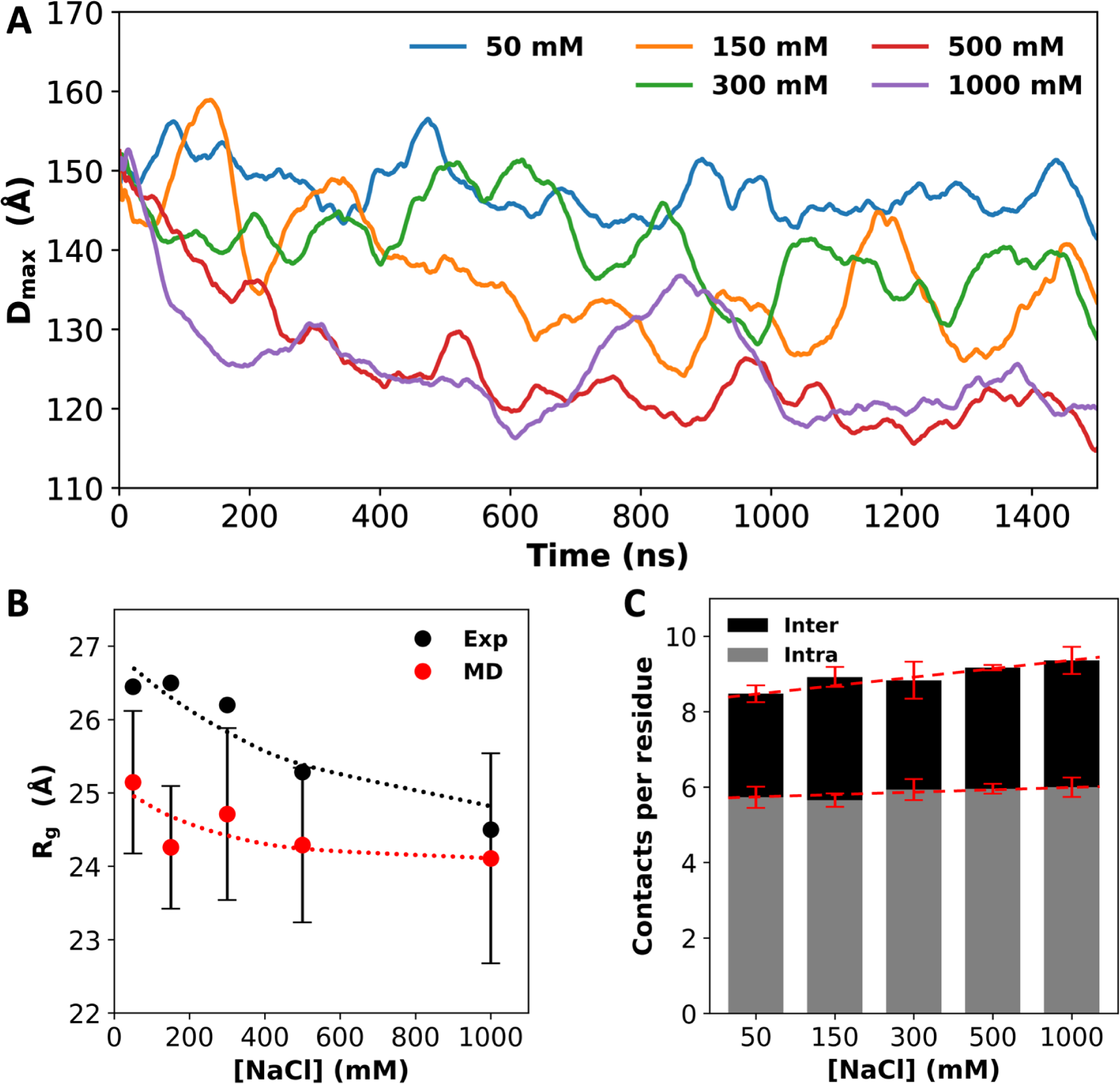
Salt effects on A1-LCD condensation and inter-chain interactions. (A) *D*_max_ values, averaged over four replicates, as a function of simulation time. (B) Radii of gyration (*R*_g_) from small-angle X-ray scattering^17^ and from MD simulations. Dotted curves are drawn to guide the eye. (C) Average number of inter- or intra-chain contacts per residue at each salt concentration. Dashed lines are drawn to show trends. Error bars represent standard deviations among four replicate simulations.

### Ions selectively bind to backbone and sidechain sites to neutralize the net charge of A1-LCD

The large net charge of A1-LCD implies significant electrostatic repulsion between chains. Potentially salt ions can neutralize the net charge. To investigate ion-protein binding, we calculated radial distribution functions (RDFs) of Cl^−^ and Na^+^ around polar groups of A1-LCD. At 1000 mM NaCl, the RDFs of Cl^−^ show a strong 1^st^ peak at 3.2 Å around Arg and Lys sidechain nitrogens and a moderate 1^st^ peak around the Gln and Asn sidechain nitrogens and the Ser sidechain oxygen (Supplementary Figure 2A). Each Arg sidechain often coordinates two Cl^−^ ions simultaneously, but each Lys sidechain coordinates only one Cl^−^ ion. A 2^nd^ peak, at ∼5 Å, is also observed around Arg and Lys sidechain nitrogens. Of the remaining polar groups, only the Cl^−^ RDFs around the Tyr sidechain oxygen and the backbone nitrogen show a weak 1^st^ peak (Supplementary Figure 2B). We used cutoffs of 4 and 6.4 Å, respectively, to define 1^st^-shell and 2^nd^-shell Cl^−^ binding. On a per-residue basis, Arg and Lys sidechains coordinate the most 1^st^-shell Cl^−^ ions, reaching ∼0.25 ions per residue (Figure 3A, left panel). In comparison, each Gln, Asn, or Ser residue, on average, coordinates ∼0.05 Cl^−^ ions. Given the large numbers of Arg and Ser residues in the A1-LCD sequence, these two residue types coordinate the most 1^st^-shell Cl^−^ ions, 21.5 and 8.4, respectively, in the 8-chain system at 1000 mM NaCl. Although the RDF around backbone nitrogens has only a weak 1^st^ peak, given their large number (131 per chain), they actually coordinate 13.4, or the second most 1^st^-shell Cl^−^ ions. Among these 13.4 Cl^−^ ions, 55% are coordinated with Gly residues. A large part of this high percentage is due to the enrichment of Gly (39% of all residues) in the A1-LCD sequence, but Gly is additionally favored for its lack of sidechain, which allows the close approach of ions to the backbone.

**Figure 3.**
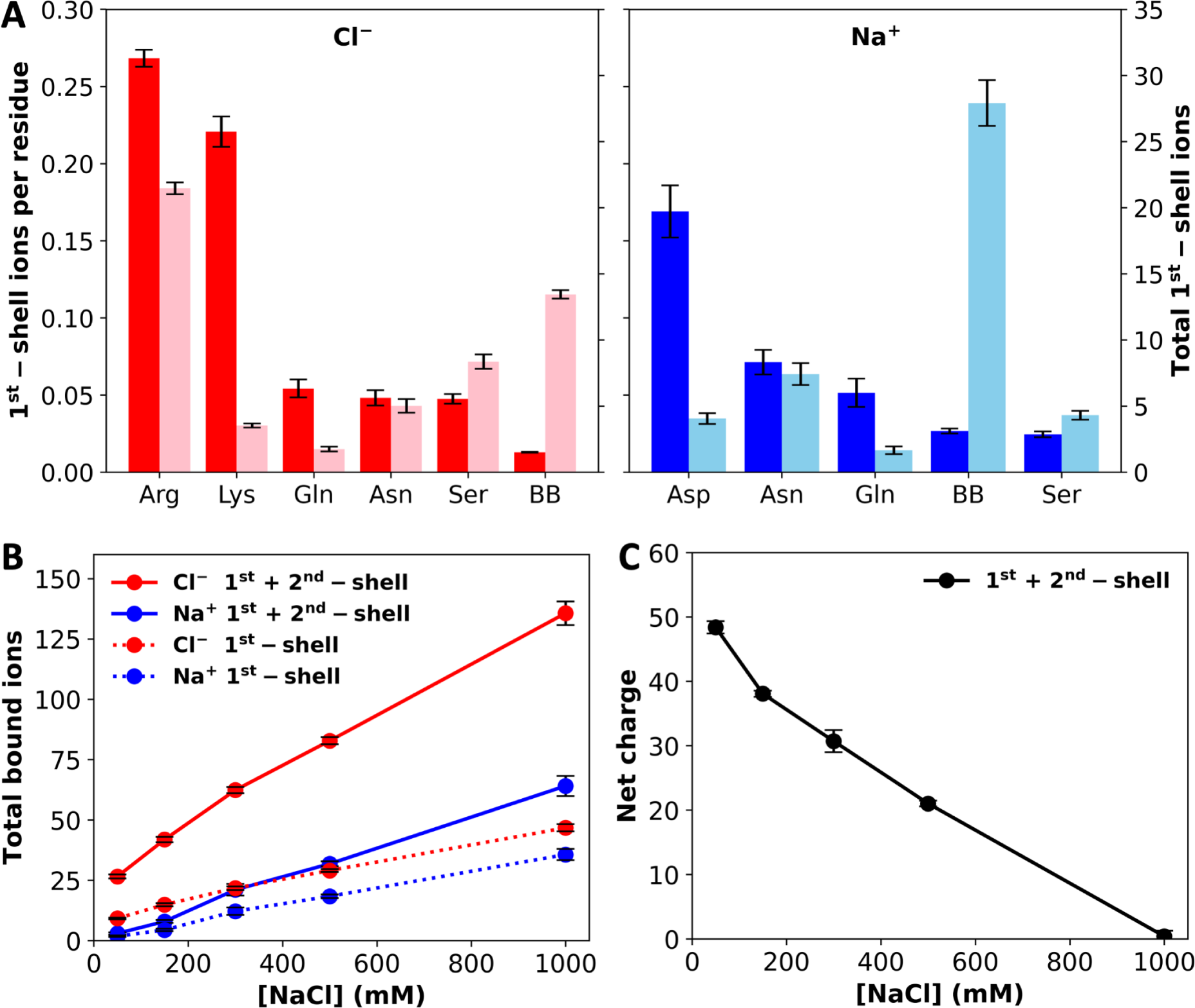
Levels of ion binding at different NaCl concentrations. (A) Number of 1^st^-shell ions, with (darker colors; tick marks on the left vertical axis) and without (lighter colors; tick marks on the right vertical axis) normalization by the number of residues of a given amino-acid type, at 1000 mM NaCl. (B) Total number of 1^st^-shell only or 1^st^- and 2^nd^-shell ions. (C) Net charge of the system with 1^st^- and 2^nd^-shell ions included. Error bars represent standard deviations among four replicate simulations.

Na^+^ shows a very strong preference for the Asp carboxyl in 1^st^-shell coordination, with a peak RDF value of 9.2 at 2.2 Å (Supplementary Figure 3A). Each Asp sidechain carboxyl typically coordinates a single Na^+^ ion, usually in a bifurcated geometry. In addition, Na^+^ shows a strong preference in 1^st^-shell coordination with Asn and Gln sidechain amide oxygens and the backbone carbonyl oxygen as well as a moderate preference with the Ser sidechain oxygen. A 2^nd^ peak, at ∼4.2 Å, is also observed in the Na^+^ RDF around the Asp carboxyl. Of the remaining polar groups, only a weak 1^st^ peak is seen in the RDF around the Tyr sidechain oxygen (Supplementary Figure 3B). We used cutoffs of 3 and 5.4 Å, respectively, to define 1^st^-shell and 2^nd^-shell Na^+^ binding. On a per-residue basis, the Asp sidechain coordinates the most 1^st^-shell Na^+^ ions, reaching 0.17 ions per residue (Figure 3A, right panel). The per-residue Na^+^ ion number reduces to ∼0.06 for Asn and Gln sidechain oxygens and further to ∼0.03 for the backbone oxygen and Ser sidechain oxygen. Again, due to their large number, backbone oxygens coordinate a very large number, 27.9, of 1^st^-shell Na^+^ in the 8-chain system at 1000 mM NaCl. This number dwarfs the counterparts for sidechains, the largest of which are 7.4, 4.1, and 4.3 respectively, for Asn, Asp, and Ser. For coordination with the backbone, similar to the case with Cl^−^, 55% of the 27.9 Na^+^ ions involve Gly residues, again reflecting the fact that this amino acid allows the close approach of ions to the backbone. On the other hand, the largest number of 1^st^-shell Cl^−^ ions are coordinated with Arg sidechains but the largest number of 1^st^-shell Na^+^ ions are coordinated with backbone carbonyls. Whereas each Arg sidechain often coordinates two Cl^−^ ions, multiple backbone carbonyls often coordinate a single Na^+^ ion, with at least two backbone carbonyl partners for 30.6 of the 64.1 bound Na^+^ ions (1^st^- and 2^nd^-shell).

In Figure 3B, we display the total numbers of 1^st^-shell Cl^−^ and Na^+^ ions in the 8-chain systems as a function of NaCl concentration. Both the 1^st^-shell Cl^−^ and Na^+^ ions increase with increasing NaCl concentration, and Cl^−^ ions always outnumber Na^+^ ions. The difference remains almost constant, with around 10 more 1^st^-shell Cl^−^ ions than 1^st^-shell Na^+^ ions. This difference is not enough to neutralize the total charge, +72, on the protein chains. However, when 2^nd^-shell ions are also included, the excess of bound Cl^−^ ions over bound Na^+^ ions grows quickly with increasing NaCl concentration. Correspondingly, the net charge of the system reduces to zero at 1000 mM NaCl (Figure 3C). Net charge repulsion explains why the chains maintain their initial loose configuration in the simulations at 50 mM NaCl (Figure 1C) and the observed absence of phase separation at this low salt concentration.^17^ At high salt, the protein net charge is completely neutralized by 1^st^- and 2^nd^-shell ions; hence the protein chains condense (Figure 1D) and phase separation was readily observed.^17^

### Ions act as bridges between protein chains to drive condensation

In addition to charge neutralization, we suspected that ions could also fortify intermolecular interactions by bridging between proteins, similar to the role played by ATP molecules in driving phase separation of positively charged IDPs.^36^ Indeed, we found that Cl^−^ has a tendency to bind with Arg and other sidechains from multiple chains and likewise, Na^+^ has a tendency to bind with backbone carbonyls and sidechain oxygens from multiple chains (Figure 4A).

**Figure 4.**
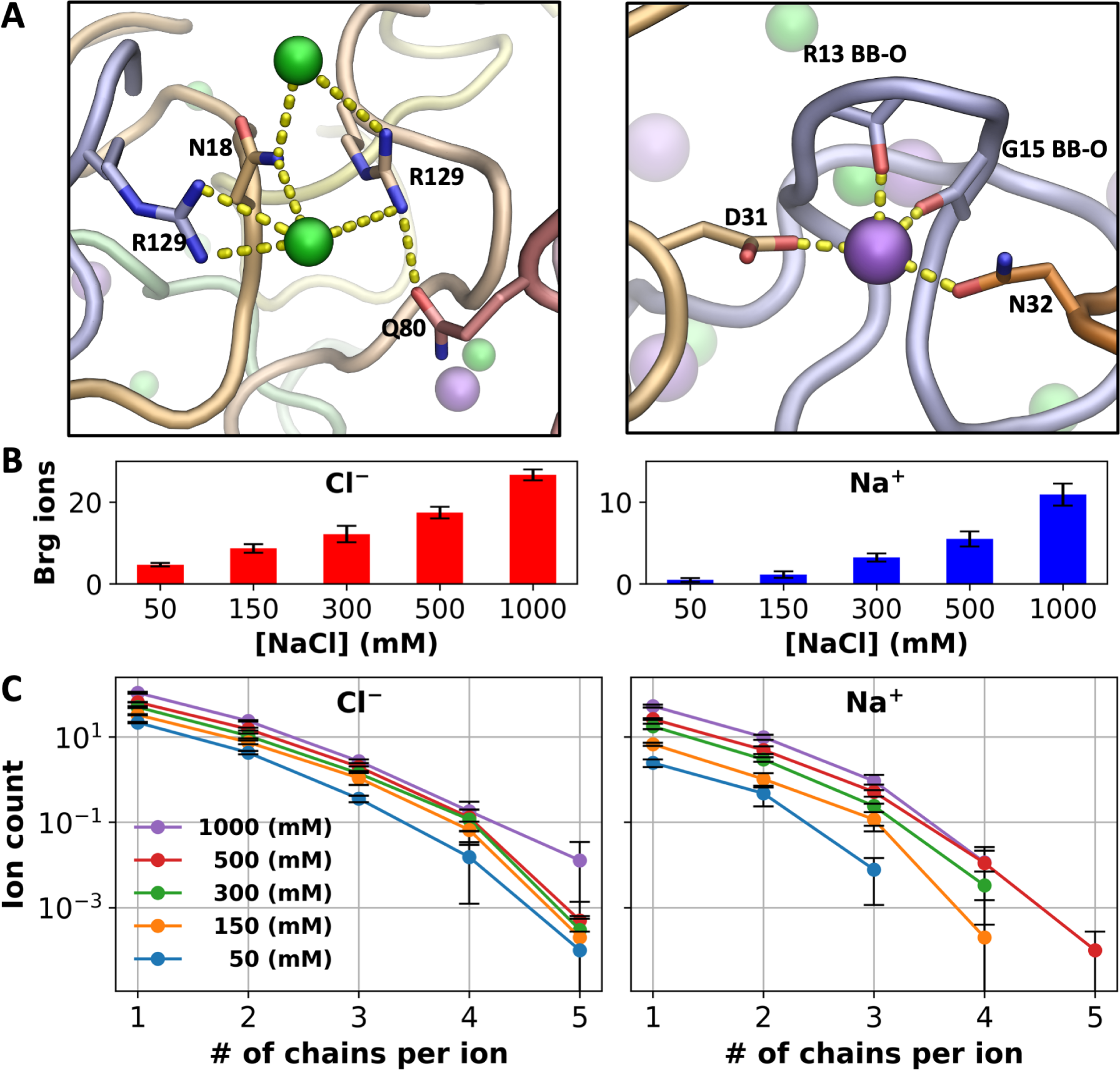
Bridging ions. (A) Examples of chain bridging by Cl^−^ and Na^+^; Cl^-^ ions are coordinated to Arg and other sidechains whereas Na^+^ ions are coordinated to both backbone carbonyls (including from Gly) and sidechain oxygens. (B) Average number of Cl^−^ or Na^+^ ions engaged in bridging between A1-LCD chains. (C) Average number of ions bound in 1^st^- and 2^nd^-shell sites lined by a given number of A1-LCD chains. Error bars represent standard deviations among four replicate simulations.

To quantify this tendency, we calculated the number of bridging Cl^−^ or Na^+^ ions, i.e., those that bind (both 1^st^- and 2^nd^-shell) to residues on more than one A1-LCD chain (Figure 4B). Close to 20% of all 1^st^- and 2^nd^-shell ions bridge between A1-LCD chains. For both Cl^−^ and Na^+^, the average number of bridging ions increases with increasing salt, but the pace of increase is much greater for Na^+^ than for Cl^−^, commensurate with the trend shown by the total number of bound ions of each type (Figure 3B). The number of bridging ions is 4.7 for Cl^−^ and only 0.5 for Na^+^ in the 8-chain system at 50 mM NaCl, and increases to 26.6 and 10.9, respectively, at 1000 mM NaCl. The greater fold change of bridging Na^+^ ions (23-fold, vs 6-fold for Cl^−^) is also apparent when we break the bridging ions according to the number of A1-LCD chains being bridged (Figure 4C). For Cl^−^, the curves plotting the ion count against the number of bridged chains are close to each other among the different NaCl concentrations, but the counterparts for Na^+^ are more spread out.

These disparate effects of salt concentration on chain bridging by Cl^−^ and Na^+^ can be explained by the difference in coordination properties between the two ion types presented above. Cl^−^ strongly prefers Arg sidechains and, even at low salt, occupies a large number of sites lined by them, of which ∼20% are bridging sites. However, there is a relatively limited supply of Arg sidechains (a total of 80 in the 8-chain system) and hence the fold change in bridging Cl^−^ ions upon salt increase is somewhat tempered. In contrast, although Na^+^ prefers Asp sidechains, there are very few of those in the system; instead, Na^+^ predominantly binds to backbone carbonyls. As the latter binding is relatively weak, the number of bridging Na^+^ ions is very small at low salt. However, since there is a large supply of backbone carbonyls (a total of 1048 in the 8-chain system), at high salt, Na^+^ ions bind to a portion of these backbone carbonyls and bridge A1-LCD chains. Of the total 10.9 Na^+^ bridging ions, 6.4 do so through backbone carbonyls of at least two chains. Consequently, at low salt, chain bridging is dominated by Cl^−^ (Na^+^ only 9% of bridging ions), but at high salt, Na^+^ becomes more even (30% of bridging ions) with Cl^−^ in bridging A1-LCD chains. An important reason for the latter is Na^+^ coordination by backbone carbonyls, especially those of Gly residues (Figure 4A).

#### Salt also contributes to condensation indirectly by strengthening π-type interactions

As noted above, the number of inter-chain interactions increases by 23% when the NaCl concentration increases from 50 mM to 1000 mM. While this result could be accounted for by the two direct salt effects presented so far, i.e., charge neutralization and chain bridging, which act to condense A1-LCD, there are additional factors. A breakdown of inter-chain sidechain interactions into different types reveals that when the salt concentration is increased from 50 to 1000 mM, the number of salt bridges per chain remains nearly constant, while the numbers of cation-π, π-π, and amino-π interactions increase by 17%, 26%, and 39%, respectively (Figure 5A). That is, as A1-LCD is condensed at high salt, there is an overall increase in inter-chain interactions, but this increase is limited to π-types of interactions and excludes salt bridges. At increasing salt, more π-types of interactions are formed while no new salt bridges are formed, suggesting a strengthening of the former interactions.

**Figure 5.**
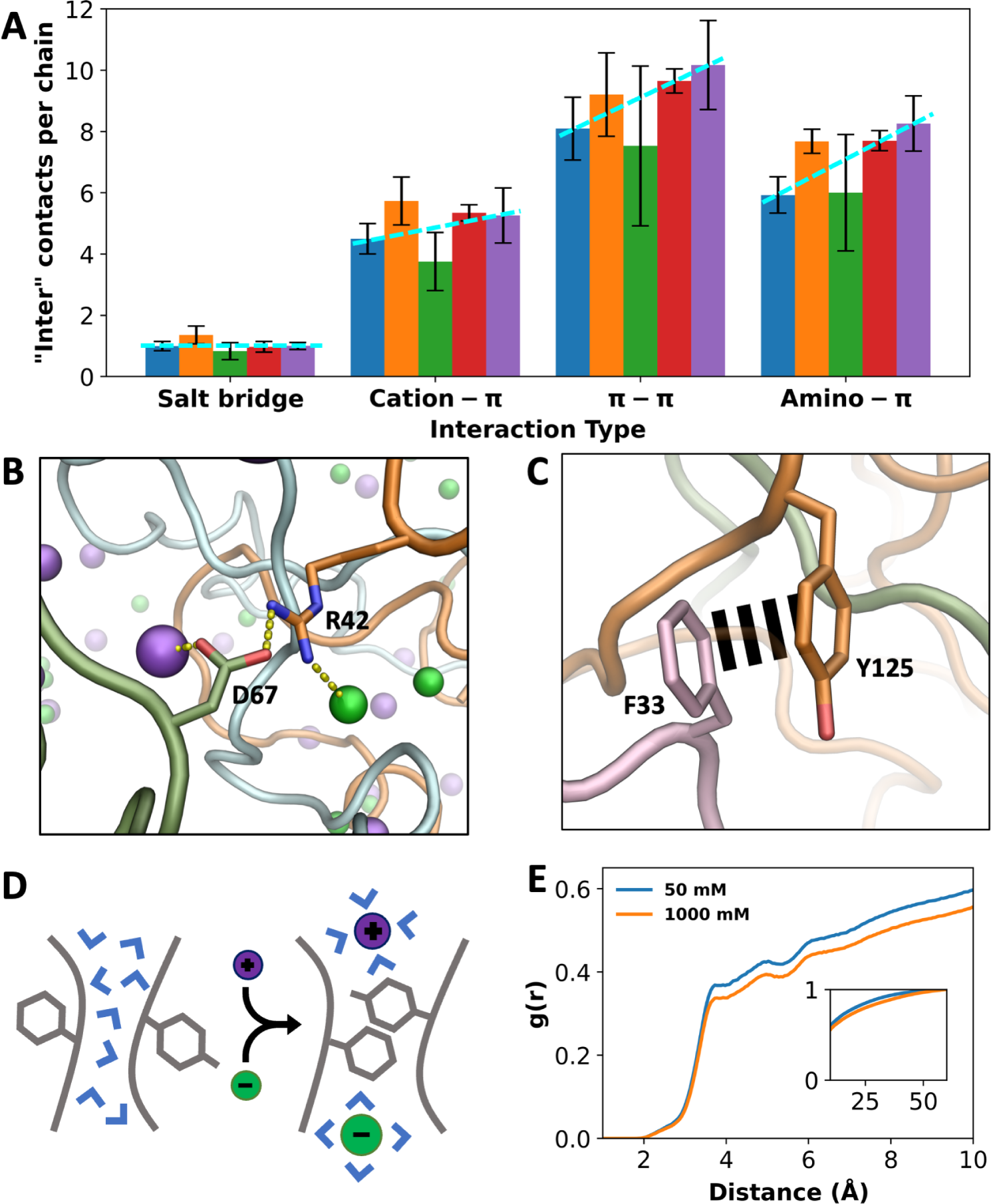
Indirect effects of ions on different types of interactions. (A) Number of inter-chain contacts per chain for each interaction type. Bars from left to right correspond to increasing salt concentrations (50, 150, 300, 500, and 1000 mM). Dashed lines are drawn to show trends. Error bars represent standard deviations among four replicate simulations. (B) An inter-chain salt bridge, with ion coordination by the partner sidechains. (C) An inter-chain π-π interaction, free of ion coordination. (D) Schematic showing a π-π interaction facilitated by high salt, via drawing water away from the interaction partners. (E) Radial distribution functions of water around Tyr residues that interact with Phe, Arg, Lys, Gln, and Asn. Lower values at high salt demonstrate water withdrawal. Inset shows radial distribution functions approaching 1.

The apparent null effect of salt on salt bridge formation can be attributed to the canceling of two opposing effects: chain condensation by the direct effects of salt may potentially shorten the distances between salt-bridge partners and thereby strengthen salt bridges but competition of ions with the salt-bridge partners for coordination (Supplementary Figures 2A and 3A) may potentially weaken salt bridges. Indeed, at 1000 mM NaCl, cationic and anionic partners in a salt bridge are often found to also coordinate Cl^−^ and Na^+^, respectively (Figure 5B). In comparison, one partner in cation-π interactions and both partners in π-π and amino-π interactions have limited abilities to coordinate with ions, so the effects from ion competition are tempered.

For π-types of interactions, instead of a weakening mechanism through ion competition, salt may exert a strengthening effect by drawing water away from the interaction partners (Figure 5D). We calculated the RDFs of water around Tyr sidechains that form π-types of interactions and found decreases in water density when the salt concentration is increased from 50 to 1000 mM (Figure 5E and Supplementary Figure 4A-C). The decreases in water density follow the order cation-π < π-π < amino-π, which is the same order as the corresponding increases in these three types of interactions. As a control, no decrease in water density is seen around Asp sidechains that form salt bridges (Supplementary Figure 4D). Water can interfere with and weaken π-types of interactions; by drawing water away from the interaction partners, high salt strengthens these interactions and thereby indirectly contributes to condensation. Recently the withdrawal of water from π-types of interaction partners in FUS-LCD condensates has been directly demonstrated by Raman spectroscopy.^40^

## Discussion

We have shown in atomistic detail the actions of ions in the condensation of A1-LCD over a wide range of NaCl concentrations. The MD simulations reveal that NaCl has both direct and indirect effects in driving phase separation. The first direct effect is to neutralize the net charge of A1-LCD and thereby attenuate net charge repulsion. The second direct effect is to bridge between protein chains and thereby fortify intermolecular interaction networks. In addition, high salt strengthens π-types of interactions by drawing water away from the interaction partners, thereby also indirectly driving phase separation. The net result is that, while phase separation of A1-LCD is prevented by net charge repulsion at low salt, it is enabled at intermediate salt through charge neutralization and chain bridging by ions. The drive for phase separation becomes even stronger at high salt, where π-types of interactions are strengthened.

These findings broaden our understanding of the roles of charges and ions in phase separation. That high net charge is a suppressive factor is highlighted by the strong effect of A1-LCD charge mutations on *C*_sat_.^31^ Here we have shown that this suppressive action can be countered by the addition of salt, which exerts both direct and indirect effects. One of these direct effects, i.e., neutralization of net charge, can be treated by a Debye-Hückel potential in coarse-grained simulations.^32^ However, our all-atom explicit-solvent simulations have revealed not only an additional direct effect, i.e., bridging between protein chains, but also an indirect effect, i.e., strengthening of π-type interactions. In lattice Monte Carlo simulations of a system of charged homopolymer chains plus counterions with Coulomb interactions between all charges, counterions were found to occupy sites between chains and produce effective chain-chain attraction;^41^ this effect might be viewed as a primitive form of bridging. Krainer et al.^28^ observed a reentrant salt effect on the phase separation of IDPs, and attributed the reemergence of phase separation at high salt to strengthened π-type and hydrophobic interactions that overcompensate weakened electrostatic attraction. Their conclusion was based on the salt dependences of the potentials of mean force calculated for pairs of sidechains of various types. Instead of a single pair of sidechains, our simulations are on multiple copies of protein chains. We show that, at high salt, the number of salt bridges remains constant while the numbers of π-types of interactions are elevated in the multi-chain system. Our simulations further reveal that high salt achieves the latter effect by drawing water away from π-interaction partners.

The present work puts us in a position to paint a unified picture of salt dependences of homotypic phase separation (Figure 6A, B and Table 1). The salt dependence that is most often reported is screening, where salts weaken the electrostatic attraction between protein chains and thereby suppress phase separation.^10–18^ The reentrant salt dependence reported by Krainer et al.^28^ can be seen as an extension of the screening scenario, whereby high salt overcompensates the screening effect by strengthening π-type and hydrophobic interactions and leads to reemergence of phase separation. The high net charge scenario represented by A1-LCD is similar to the reentrant scenario at high salt but differs from it at low salt, where phase separation is prevented by net charge repulsion.^17,20,22–27,29^ In this class of salt dependence, a certain amount of salt is required to start phase separation. For A1-LCD, this minimum salt is ∼100 mM NaCl.^17^ The recombinant mussel foot protein-1 (RMFP-1) presents an extreme example, which has a net charge of +24 over a sequence length of 121 and requires up to 700 mM NaCl to begin phase separation.^22^ The fourth class of salt dependence is a variation of the preceding one; here the net charge is low and no significant net charge repulsion is expected, so the protein can phase separate even without salt. For example, FUS-LCD has a low net charge of -2 and phase separates without salt.^21^ The difference between the high and low net charge scenarios is well captured by the phase-separation behaviors of TDP43-LCD at two pHs.^24^ At pH 7, this protein has a relatively low net charge of +5 and phase separates without salt; the salt dependence thus belongs to the low net charge class. When pH is lowered to 4, the six His residues in the purification tag become positively charged, raising the net charge to +11; now phase separation requires 300 mM NaCl and the salt dependence switches to the high net charge class.

**Figure 6.**
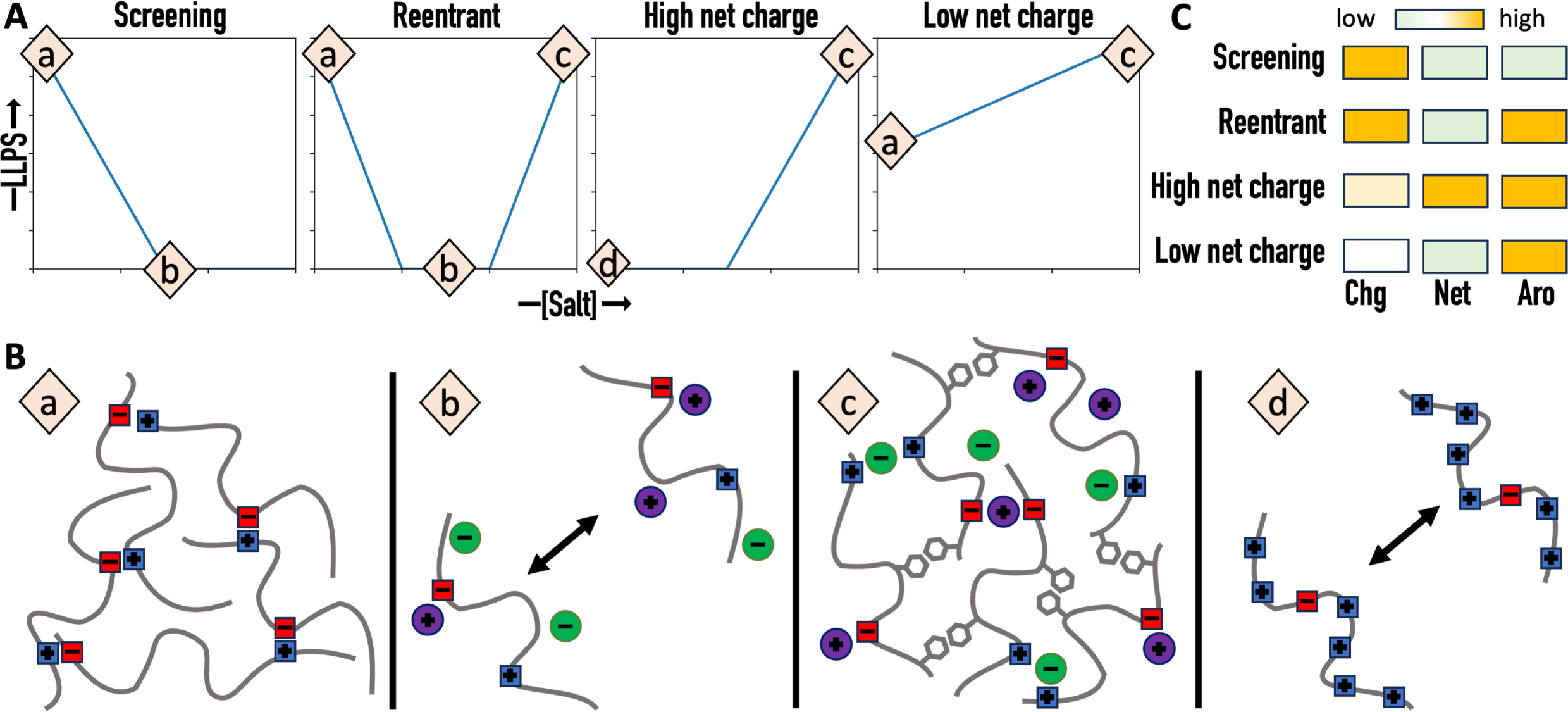
Four classes of salt dependence and their prediction from amino-acid composition. (A) Salt dependences of liquid-liquid phase separation (LLPS). (B) Charge-charge and π-type interactions and their regulation by salt. (a) Significant Charge-charge attraction. (b) Screening of charge-charge attraction by salt. (c) Strengthening of π-type interactions by high salt. (d) Repulsion due to high net charge. (C) Distinctions of the four classes of salt dependence by three determinants: charged content (“Chg”), net charge (“Net”), and aromatic content (“Aro”).

**Table 1.**
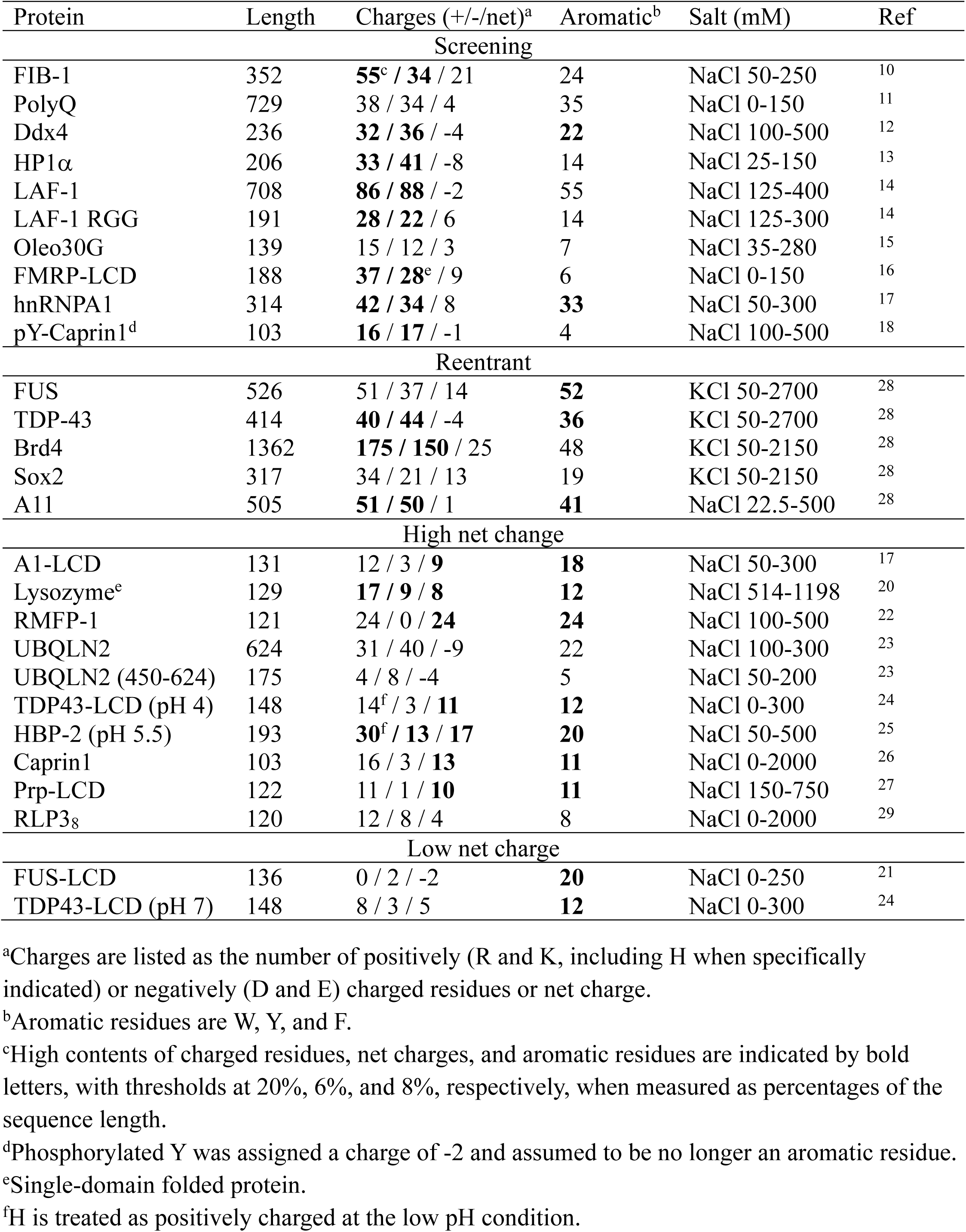
Correlation between class of salt dependence and amino-acid composition.

We can not only rationalize the four distinct classes of salt dependence but also start to predict them from the amino-acid composition of the protein (Figure 6C and Table 1). The foregoing physical understanding suggests three determinants of salt dependence class: (1) the total number of charged residues, which determines the contribution of electrostatic interactions to the drive for phase separation and the importance of salt screening; (2) the net charge, which determines the magnitude of net charge repulsion; and (3) the total number of aromatic residues, which determines whether phase separation reemerges at high salt. We predict that the screening class occurs when the charged content is high but the net charge and aromatic content are low (Figure 6C). The other three classes of salt dependence all require a high aromatic content. For the reentrant class, the increase in aromatic content is the only difference from the screening class. For the high net charge class, the net charge obviously has to be high, but that could also mean at least a moderately high charged content. Finally the low net charge class is predicted when the net charge is low but no bias is required of the charged content. The amino-acid compositional data in Table 1 validate these predictions. For example, of the 17 proteins in the classes of salt dependence that call for a high aromatic content, 12 (or 71%) actually have this feature (aromatic content > an 8% threshold). In contrast, of the 10 proteins in the class of salt dependence (i.e., “screening”) that calls for a low aromatic content, only 2 (or 20%) have an aromatic content above the threshold. In the latter two cases it remains to be tested whether phase separation would reemerge at much higher salt (and thereby resulting in a reclassification to reentrant). Likewise, of the 10 proteins in the class of salt dependence (i.e., “high net charge”) that calls for a high net charge, 7 (or 70%) actually have this feature (net charge > a 6% threshold). In contrast, of the 17 proteins in the classes of salt dependence that call for a low net charge, none has a net charge above the threshold. Lastly, of the 15 proteins in the classes of salt dependence that call for a high charge content, 11 (or 73%) actually have this feature (charged content > a 20% threshold). In comparison, of the 12 proteins in the classes of salt dependence that do not require a high charge content, only 2 (or 17%) have a charge content above the threshold.

Two of the proteins included in Table 1 are Caprin1 and its phosphorylated variant pY-Caprin1. Caprin1’s phase separation required 500 mM NaCl and was promoted by higher salt,^26^ and thus behaved as if in the high net charge class of salt dependence (Figure 6A, third panel). In contrast, pY-Caprin1’s phase separation was suppressed by salt,^18^ behaving as if in the screening class (Figure 6A, first panel). Our amino-acid composition-based predictor correctly places both proteins in these respective classes (Table 1), with Caprin1’s sequence possessing both a high net charge and a high aromatic content whereas pY-Caprin1’s sequence possessing a high level of charged residues. The disparate salt dependences of these two proteins were also correctly captured by both analytical theory and coarse-grained simulations.^18^ The simulations also showed the dominance of Cl^−^ over Na^+^ in partitioning into the dense phase of Caprin1, similar to our all-atom simulations of A1-LCD (Figure 3B); both Caprin1 and A1-LCD have a high net positive charge.

Comparing and contrasting coarse-grained simulations and all-atom simulations highlight the pros and cons of the two approaches. Coarse-grained simulations can readily capture the phase-separation process and represent the equilibrium between the dense and dilute phases. For short peptides, all-atom simulations have been shown to accomplish both of these tasks.^37^ For the mixture of ATP and a 33-residue IDP, all-atom simulations were able to capture the condensation process but not the equilibration of the resulting dense phase with a dilute phase.^36^ In this work, although the total number of atoms in the A1-LCD system, ∼half a million, is very high from a computational standpoint, the number of protein chains (i.e., 8) is still too low for a proper representation of a homogenous solution at low salt and a full equilibration between two phases at intermediate and high salt. Nevertheless, our simulations do not show condensation at low salt, thus mimicking a homogenous solution; the simulations at intermediate and high salt show condensation and thus model the dense phase. We are thus able to compare the salt effects in a homogenous solution (50 mM NaCl) and in the dense phase after phase separation (150-1000 mM NaCl) and compare the salt effects in dense phases with intermediate (150 and 300 mM NaCl) and high condensation (500 and 1000 mM NaCl). Importantly, the all-atom simulations are able to capture unusual physical effects that could be missed in coarse-grained simulations, such as strengthening π-types of interactions by drawing water away from the interaction partners. The latter effect is now confirmed by Raman spectroscopy.^40^

To conclude, salts regulate intermolecular interactions in biomolecular condensates in a variety of ways, but the net dependence of homotypic phase separation on salt concentration can be placed into four distinct classes (Figure 6A, B). Moreover, these classes are predictable from the protein amino-acid composition. Salts also exert significant effects on heterotypic condensates;^8,19^ the conclusions drawn here on homotypic condensates may prove instructive for heterotypic condensates. The fact that different IDPs respond to salts differently raises interesting questions on phase separation inside cells. For example, a fluctuation in intracellular salt concentration may promote the phase separation of some IDPs and suppress the phase separation of other IDPs. If two IDPs co-phase separate or form heterotypic condensates, salt fluctuations may change the protein composition of condensates and the relative contributions of the individual proteins to phase separation. Lastly, we note that the atomistic details of ion coordination to A1-LCD backbone or sidechain groups are reminiscent of those observed in other systems, in particular channels and transporters for Na^+^ and Cl^−^.^42–44^

### Computational Methods

#### Molecular dynamics simulations

MD simulations were performed using AMBER 18^45^ with ff14SB force field^46^ for the protein and TIP4P-D for water.^47^ The initial configuration of the 8-chain system was constructed using A1-LCD conformations from previous single-copy simulations.^39^ Specifically, 4 copies with different conformations were placed in a rectangular box, positioned with minimal inter-chain contacts within 3.5 Å. This 4-copy subsystem was duplicated to form the initial configuration of the 8-chain system (Figure 1B). The protein chains were solvated in a box with dimensions of 182 Å ξ 146 Å ξ 164 Å. For each desired salt concentration (50, 150, 300, 500, or 1000 mM), an appropriate number of water molecules were randomly selected and replaced with Cl^−^ and Na^+^ ions; excess Cl^−^ was added to neutralize the system. The total number of atoms was ∼500,000 for the 8-chain system at each salt concentration.

After energy minimization in sander (2000 steps of steepest descent and 3000 steps of conjugate gradient), each system was heated to 300 K over 100 ps with a 1 fs timestep, under constant NVT using the Langevin thermostat^48^ with a 3.0 ps^-1^ damping constant. The simulation was then continued in four replicates at constant NPT for 1.5 µs with a 2 fs timestep. The final dimensions were stabilized at ∼ 174 Å x 140 Å x 157 Å. Pressure was regulated using the Berendsen barostat^49^ with a coupling constant of 2.0 ps. All simulations were run on GPUs using *pmemd.cuda*.^50^ Bond lengths involving hydrogens were constrained using the SHAKE algorithm.^51^ Long-range electrostatic interactions were treated by the particle mesh Ewald method.^52^ A cutoff distance of 10 Å was used for the nonbonded interactions. Frames were saved every 200 ps and the last 500 ns was used for analysis.

#### Data analysis

*D*_max_ was calculated using the minmax function in VMD.^53^ The differences between the maximum and minimum values of *x*, *y*, and *z* coordinates of all protein atoms were obtained; the largest of the three difference values was taken as *D*_max_. To avoid issues with periodic boundary conditions, this calculation was repeated 8 times, each time the simulation box was centered on one of the A1-LCD chains. The smallest of the 8 *D*_max_ values is reported in Figure 2A. Radius of gyration (*R*_g_) for each chain was calculated using the radgyr function in CPPTRAJ and then averaged over the 8 chains in each simulation (Figure 2B).^54^ For inter- and intrachain contacts per residue (Figure 2C), the number of other residues in contact with a given residue was found using the distance mask function in CPPTRAJ with a cutoff of 6 Å between heavy atoms. Again, to avoid issues with the periodic boundary conditions, each chain was centered individually before inter- and intrachain contacts were calculated for each residue within that chain. In the case of intrachain contacts, the two nearest neighboring residues of a given residue were excluded as contact partners. Interchain contacts per chain (Figure 5A) were found in a similar way using only sidechain heavy atoms, and were broken into interaction types based on the residue types of the interaction partners. Interaction types were defined as follows: salt bridges were between Arg/Lys and Asp, cation-π interactions were between Arg/Lys and Tyr/Phe, amino-π interactions were between Asn/Gln and Tyr/Phe, and π-π interactions were between a Tyr/Phe and another Tyr/Phe. To characterize chain reconfiguration, the RMSFs of the chains were calculated separately and then averaged over the 8 chains (Supplementary Figure 1).

Radial distribution functions (RDFs) for ions around protein atoms were calculated in CPPTRAJ using the radial command with a bin size of 0.05 Å and a specific selection of the protein atom type for either Cl^−^or Na^+^ at the center. These RDF plots were used to select cutoffs for 1^st^- and 2^nd^-shell ion binding; the cutoffs were 4.0 and 6.4 Å, respectively, for Cl^−^ and 3.0 and 5.4 Å, respectively, for Na^+^ (Supplementary Figures 2 and 3). The number of 1^st^-shell or 1^st^- and 2^nd^-shell ions interacting with N and O atoms in a given sidechain type, the backbone, or the 8 chains together (Figure 3) was calculated using CPPTRAJ, by specifying a protein selection, an ion selection (all Cl^−^ ions or all Na^+^ ions), and a cutoff distance. Protein chains were centered one at a time and each time, the protein selection was limited to the central chain. The resulting ion-protein interactions for the 8 chains were then combined and custom python scripts were used to collect unique ions in the total ion count for each protein selection (Figure 3A, right ordinate; Figure 3B). For ions interacting with a given sidechain type, the total ion count was normalized by the number of residues of that type to get the ion count per residue (Figure 3A, left ordinate). The net charge of the system was calculated by taking the total charge of the 8 protein chains (+72) and adding and subtracting, respectively, the numbers of bound Na^+^ and Cl^-^ ions within the 2^nd^-shell cutoffs (Figure 3C). The total bound ions in Figure 3B were further divided according to the number of chains with which each ion was interacting in the same frame. Ions were considered bridging if they interacted with two or more chains (based on 2^nd^-shell cutoffs) in the same frame (Figure 4B). Those interacting with an exact number of chains (e.g., 3) were identified when the same ion was found in the interaction lists of that many chains (Figure 4C). Error bars in all graphs represent the standard deviations among the four replicate simulations at each salt concentration.

RDFs for water molecules around Tyr residues were calculated using VMD with a bin size of 0.05 Å (Figure 5E and Supplementary Figure 4A-C). The water atom selection was all its three atoms and the Tyr atom selection was its 6-carbon ring. Moreover, only Tyr residues that interact with a specific residue type (within 6 Å of the 6-carbon ring) were selected: the partner residues were Phe (6-carbon ring; for π-π), Arg and Lys (sidechain N atoms; for cation-π), or Gln and Asn (sidechain N atoms; for amino-π). As a control, RDFs for water were also calculated around Asp sidechain oxygens that interact with Arg and Lys sidechain nitrogens (Supplementary Figure 4D).

## Acknowledgment

This work was supported by National Institutes of Health Grant GM118091.

## References

(1) Mitrea, D. M.; Cika, J. A.; Stanley, C. B.; Nourse, A.; Onuchic, P. L.; Banerjee, P. R.; Phillips, A. H.; Park, C.-G.; Deniz, A. A.; Kriwacki, R. W. Self-interaction of NPM1 modulates multiple mechanisms of liquid–liquid phase separation. Nat Commun 2018, 9, 842.

(2) Protter, D. S. W.; Parker, R. Principles and Properties of Stress Granules. Trends Cell Biol 2016, 26, 668–679.

(3) Das, S.; Lin, Y. H.; Vernon, R. M.; Forman-Kay, J. D.; Chan, H. S. Comparative Roles of Charge, Pi, and Hydrophobic Interactions in Sequence-Dependent Phase Separation of Intrinsically Disordered Proteins. Proc Natl Acad Sci U S A 2020, 117, 28795–28805.

(4) Dignon, G. L.; Best, R. B.; Mittal, J. Biomolecular Phase Separation: From Molecular Driving Forces to Macroscopic Properties. Annu Rev Phys Chem 2020, 71, 53–75.

(5) Zhou, H. X.; Kota, D.; Qin, S.; Prasad, R. Fundamental Aspects of Phase-Separated Biomolecular Condensates. Chem Rev 2024, 124, 8550–8595.

(6) Baldwin, R. L. How Hofmeister ion interactions affect protein stability. Biophys J 1996, 71, 2056–2063.

(7) Zhou, H. X.; Pang, X. Electrostatic Interactions in Protein Structure, Folding, Binding, and Condensation. Chem Rev 2018, 118, 1691–1741.

(8) Farag, M.; Borcherds, W. M.; Bremer, A.; Mittag, T.; Pappu, R. V. Phase separation of protein mixtures is driven by the interplay of homotypic and heterotypic interactions. Nat Commun 2023, 14, 5527.

(9) Hazra, M. K.; Levy, Y. Cross-Talk of Cation−π Interactions with Electrostatic and Aromatic Interactions: A Salt-Dependent Trade-off in Biomolecular Condensates. J Phys Chem Lett 2023, 14, 8460–8469.

(10) Berry, J.; Weber, S. C.; Vaidya, N.; Haataja, M.; Brangwynne, C. P. RNA transcription modulates phase transition-driven nuclear body assembly. Proc Natl Acad Sci U S A 2015, 112, E5237–5245.

(11) Zhang, H.; Elbaum-Garfinkle, S.; Langdon, E. M.; Taylor, N.; Occhipinti, P.; Bridges, A. A.; Brangwynne, C. P.; Gladfelter, A. S. RNA Controls PolyQ Protein Phase Transitions. Mol Cell 2015, 60, 220–230.

(12) Brady, J. P.; Farber, P. J.; Sekhar, A.; Lin, Y. H.; Huang, R.; Bah, A.; Nott, T. J.; Chan, H. S.; Baldwin, A. J.; Forman-Kay, J. D.et al. Structural and hydrodynamic properties of an intrinsically disordered region of a germ cell-specific protein on phase separation. Proc Natl Acad Sci U S A 2017, 114, E8194–E8203.

(13) Strom, A. R.; Emelyanov, A. V.; Mir, M.; Fyodorov, D. V.; Darzacq, X.; Karpen, G. H. Phase separation drives heterochromatin domain formation. Nature 2017, 547, 241–245.

(14) Wei, M. T.; Elbaum-Garfinkle, S.; Holehouse, A. S.; Chen, C. C.; Feric, M.; Arnold, C. B.; Priestley, R. D.; Pappu, R. V.; Brangwynne, C. P. Phase behaviour of disordered proteins underlying low density and high permeability of liquid organelles. Nat Chem 2017, 9, 1118–1125.

(15) Reed, E. H.; Hammer, D. A. Redox sensitive protein droplets from recombinant oleosin. Soft Matter 2018, 14, 6506–6513.

(16) Tsang, B.; Arsenault, J.; Vernon, R. M.; Lin, H.; Sonenberg, N.; Wang, L.-Y.; Bah, A.; Forman-Kay, J. D. Phosphoregulated FMRP Phase Separation Models Activity-Dependent Translation through Bidirectional Control of mRNA Granule Formation. Proc. Natl. Acad. Sci. USA 2019, 116, 4218–4227.

(17) Martin, E. W.; Thomasen, F. E.; Milkovic, N. M.; Cuneo, M. J.; Grace, C. R.; Nourse, A.; Lindorff-Larsen, K.; Mittag, T. Interplay of folded domains and the disordered low-complexity domain in mediating hnRNPA1 phase separation. Nucleic Acids Res 2021, 49, 2931–2945.

(18) Lin, Y.-H.; Kim, T. H.; Das, S.; Pal, T.; Wessén, J.; Rangadurai, A. K.; Kay, L. E.; Forman-Kay, J. D.; Chan, H. S. Electrostatics of Salt-Dependent Reentrant Phase Behaviors Highlights Diverse Roles of ATP in Biomolecular Condensates. eLife 2024, DOI:10.7554/elife.100284.1 10.7554/elife.100284.1.

(19) Galvanetto, N.; Ivanović, M. T.; Chowdhury, A.; Sottini, A.; Nüesch, M. F.; Nettels, D.; Best, R. B.; Schuler, B. Extreme dynamics in a biomolecular condensate. Nature 2023, 619, 876–883.

(20) Muschol, M.; Rosenberger, F. Liquid–liquid phase separation in supersaturated lysozyme solutions and associated precipitate formation/crystallization. J Chem Phys 1997, 107, 1953–1962.

(21) Burke, K. A.; Janke, A. M.; Rhine, C. L.; Fawzi, N. L. Residue-by-Residue View of In Vitro FUS Granules that Bind the C-Terminal Domain of RNA Polymerase II. Mol Cell 2015, 60, 231–241.

(22) Kim, S.; Yoo, H. Y.; Huang, J.; Lee, Y.; Park, S.; Park, Y.; Jin, S.; Jung, Y. M.; Zeng, H.; Hwang, D. S.et al. Salt Triggers the Simple Coacervation of an Underwater Adhesive When Cations Meet Aromatic pi Electrons in Seawater. ACS Nano 2017, 11, 6764–6772.

(23) Dao, T. P.; Kolaitis, R. M.; Kim, H. J.; O’Donovan, K.; Martyniak, B.; Colicino, E.; Hehnly, H.; Taylor, J. P.; Castaneda, C. A. Ubiquitin Modulates Liquid-Liquid Phase Separation of UBQLN2 via Disruption of Multivalent Interactions. Mol Cell 2018, 69, 965–978 e966.

(24) Babinchak, W. M.; Haider, R.; Dumm, B. K.; Sarkar, P.; Surewicz, K.; Choi, J. K.; Surewicz, W. K. The role of liquid-liquid phase separation in aggregation of the TDP-43 low-complexity domain. J Biol Chem 2019, 294, 6306–6317.

(25) Le Ferrand, H.; Duchamp, M.; Gabryelczyk, B.; Cai, H.; Miserez, A. Time-Resolved Observations of Liquid-Liquid Phase Separation at the Nanoscale Using in Situ Liquid Transmission Electron Microscopy. J Am Chem Soc 2019, 141, 7202–7210.

(26) Wong, L. E.; Kim, T. H.; Muhandiram, D. R.; Forman-Kay, J. D.; Kay, L. E. NMR Experiments for Studies of Dilute and Condensed Protein Phases: Application to the Phase-Separating Protein CAPRIN1. J Am Chem Soc 2020, 142, 2471–2489.

(27) Agarwal, A.; Rai, S. K.; Avni, A.; Mukhopadhyay, S. An intrinsically disordered pathological prion variant Y145Stop converts into self-seeding amyloids via liquid-liquid phase separation. Proc Natl Acad Sci U S A 2021, 118.

(28) Krainer, G.; Welsh, T. J.; Joseph, J. A.; Espinosa, J. R.; Wittmann, S.; de Csillery, E.; Sridhar, A.; Toprakcioglu, Z.; Gudiskyte, G.; Czekalska, M. A.et al. Reentrant liquid condensate phase of proteins is stabilized by hydrophobic and non-ionic interactions. Nat Commun 2021, 12, 1085.

(29) Otis, J. B.; Sharpe, S. Sequence Context and Complex Hofmeister Salt Interactions Dictate Phase Separation Propensity of Resilin-like Polypeptides. Biomacromolecules 2022, 23, 5225–5238.

(30) Martin, E. W.; Holehouse, A. S.; Peran, I.; Farag, M.; Incicco, J. J.; Bremer, A.; Grace, C. R.; Soranno, A.; Pappu, R. V.; Mittag, T. Valence and patterning of aromatic residues determine the phase behavior of prion-like domains. Science 2020, 367, 694–699.

(31) Bremer, A.; Farag, M.; Borcherds, W. M.; Peran, I.; Martin, E. W.; Pappu, R. V.; Mittag, T. Deciphering how naturally occurring sequence features impact the phase behaviours of disordered prion-like domains. Nat Chem 2022, 14, 196–207.

(32) Tesei, G.; Lindorff-Larsen, K. Improved Predictions of Phase Behaviour of Intrinsically Disordered Proteins by Tuning the Interaction Range. Open Res Eur 2023, 2, 94.

(33) Lin, Y.-H.; Forman-Kay, J. D.; Chan, H. S. Sequence-Specific Polyampholyte Phase Separation in Membraneless Organelles. Phys Rev Lett 2016, 117, 178101.

(34) Lin, Y. H.; Brady, J. P.; Chan, H. S.; Ghosh, K. A unified analytical theory of heteropolymers for sequence-specific phase behaviors of polyelectrolytes and polyampholytes. J Chem Phys 2020, 152, 045102.

(35) Garaizar, A.; Espinosa, J. R. Salt dependent phase behavior of intrinsically disordered proteins from a coarse-grained model with explicit water and ions. J Chem Phys 2021, 155. (acccessed 6/6/2024).

(36) Kota, D.; Prasad, R.; Zhou, H. X. Adenosine Triphosphate Mediates Phase Separation of Disordered Basic Proteins by Bridging Intermolecular Interaction Networks. J Am Chem Soc 2024, 146, 1326–1336.

(37) Zhang, Y.; Prasad, R.; Su, S.; Lee, D.; Zhou, H.-X. Amino Acid-Dependent Phase Equilibrium and Material Properties of Tetrapeptide Condensates. Cell Rep Phys Sci 2024, 5, 102218.

(38) Rekhi, S.; Garcia, C. G.; Barai, M.; Rizuan, A.; Schuster, B. S.; Kiick, K. L.; Mittal, J. Expanding the molecular language of protein liquid–liquid phase separation. Nat Chem 2024, DOI:10.1038/s41557-024-01489-x 10.1038/s41557-024-01489-x.

(39) Dey, S.; MacAinsh, M.; Zhou, H. X. Sequence-Dependent Backbone Dynamics of Intrinsically Disordered Proteins. J Chem Theory Comput 2022, 18, 6310–6323.

(40) Joshi, A.; Avni, A.; Walimbe, A.; Rai, S. K.; Sarkar, S.; Mukhopadhyay, S. Hydrogen-Bonded Network of Water in Phase-Separated Biomolecular Condensates. J Phys Chem Lett 2024, 15, 7724–7734.

(41) Orkoulas, G.; Kumar, S. K.; Panagiotopoulos, A. Z. Monte carlo study of coulombic criticality in polyelectrolytes. Phys Rev Lett 2003, 90, 048303.

(42) Mancusso, R.; Gregorio, G. G.; Liu, Q.; Wang, D. N. Structure and mechanism of a bacterial sodium-dependent dicarboxylate transporter. Nature 2012, 491, 622–626.

(43) Mita, K.; Sumikama, T.; Iwamoto, M.; Matsuki, Y.; Shigemi, K.; Oiki, S. Conductance selectivity of Na(+) across the K(+) channel via Na(+) trapped in a tortuous trajectory. Proc Natl Acad Sci U S A 2021, 118.

(44) Leisle, L.; Lam, K.; Dehghani-Ghahnaviyeh, S.; Fortea, E.; Galpin, J. D.; Ahern, C. A.; Tajkhorshid, E.; Accardi, A. Backbone amides are determinants of Cl(-) selectivity in CLC ion channels. Nat Commun 2022, 13, 7508.

(45) Case, D. A.; Ben-Shalom, I. Y.; Brozell, S. R.; Cerutti, D. S.; Cheatham, T. E.; Cruzeiro, V. W. D.; Darden, T. A.; Duke, R. E.; Ghoreishi, D.; Gilson, M. K. AMBER 2018, University of California, San Francisco. 2018.

(46) Maier, J. A.; Martinez, C.; Kasavajhala, K.; Wickstrom, L.; Hauser, K. E.; Simmerling, C. ff14SB: Improving the Accuracy of Protein Side Chain and Backbone Parameters from ff99SB. J Chem Theory Comput 2015, 11, 3696–3713.

(47) Piana, S.; Donchev, A. G.; Robustelli, P.; Shaw, D. E. Water dispersion interactions strongly influence simulated structural properties of disordered protein states. J Phys Chem B 2015, 119, 5113–5123.

(48) Pastor, R. W.; Brooks, B. R.; Szabo, A. An analysis of the accuracy of langevin and molecular dynamics algorithms. Mol Phys 1988, 65, 1409–1419.

(49) Berendsen, H. J. C.; Postma, J. P. M.; van Gunsteren, W. F.; DiNola, A.; Haak, J. R. Molecular dynamics with coupling to an external bath. J Chem Phys 1984, 81, 3684–3690.

(50) Salomon-Ferrer, R.; Gotz, A. W.; Poole, D.; Le Grand, S.; Walker, R. C. Routine Microsecond Molecular Dynamics Simulations with AMBER on GPUs. 2. Explicit Solvent Particle Mesh Ewald. J Chem Theory Comput 2013, 9, 3878–3888.

(51) Ryckaert, J. P.; Ciccotti, G.; Berendsen, H. J. C. Numerical integration of the cartesian equations of motion of a system with constraints: molecular dynamics of n-alkanes. J Comput Phys 1977, 23, 327–341.

(52) Essmann, U.; Perera, L.; Berkowitz, M. L.; Darden, T.; Lee, H.; Pedersen, L. G. A smooth particle mesh Ewald method. J Chem Phys 1995, 103, 8577–8593.

(53) Humphrey, W.; Dalke, A.; Schulten, K. VMD: visual molecular dynamics. J Mol Graph 1996, 14, 33–38, 27-38.

(54) Roe, D. R.; Cheatham, T. E., 3rd. PTRAJ and CPPTRAJ: Software for Processing and Analysis of Molecular Dynamics Trajectory Data. J Chem Theory Comput 2013, 9, 3084–3095.

