## Supplementary Figures for "Direct and Indirect Salt Effects on Homotypic Phase Separation"

*Supplementary Information*

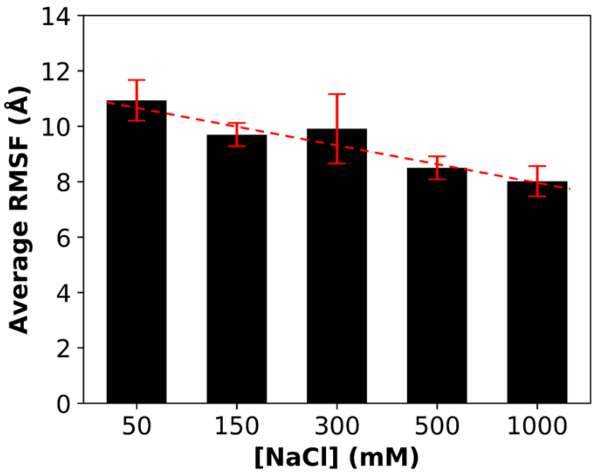


Supplementary Figure 1. Salt dependence of the average root-mean-square-fluctuation (RMSF) among the 8 chains. Error bars represent standard deviations among four replicate simulations.

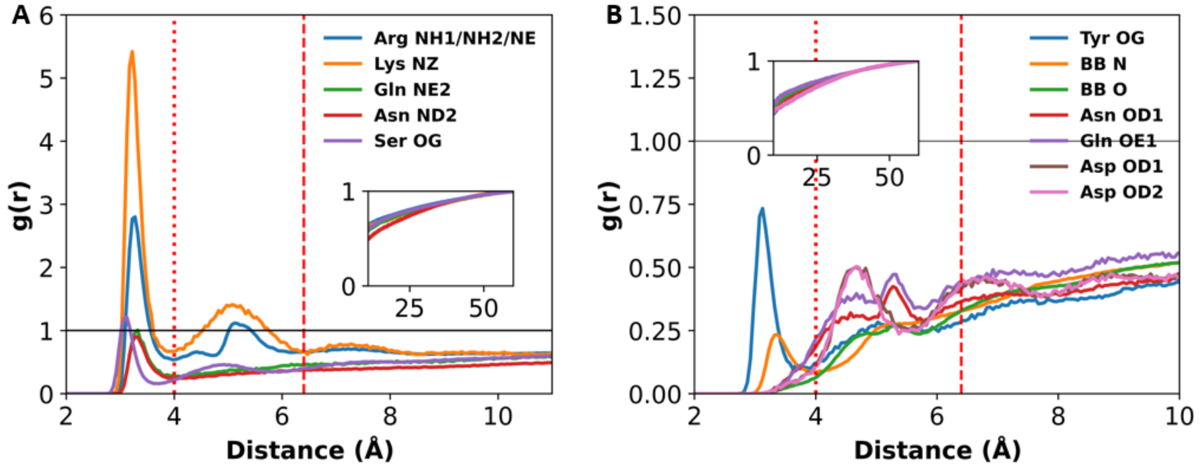


Supplementary Figure 2. Radial distributions functions of $Cl^{-}$ around sidechain and backbone N and O atoms. (A) Groups having RDF values > or close to 1. (B) Other groups. Two vertical lines indicate cutoff distances for 1st- and 2nd-shell coordination. Insets show the approach of radial distribution functions to 1.

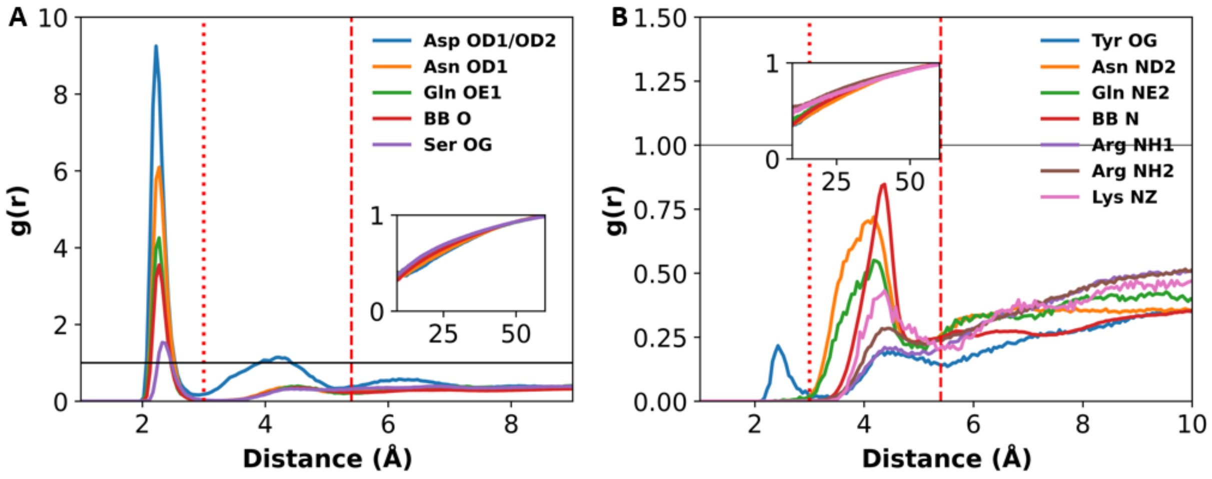


Supplementary Figure 3. Radial distributions functions of $Na^+$ around sidechain and backbone O and N atoms. (A) Groups having RDF values > 1. (B) Other groups. Two vertical lines indicate cutoff distances for 1$^{st}$- and 2$^{nd}$-shell coordination. Insets show the approach of radial distribution functions to 1.

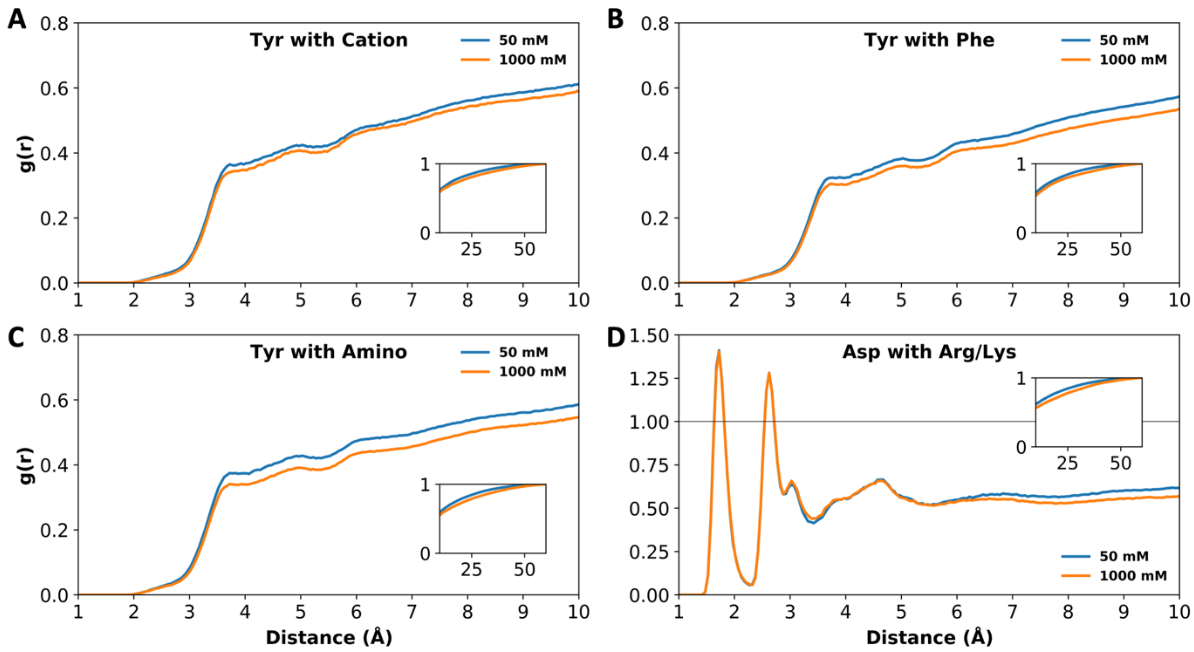


Supplementary Figure 4. Radial distribution functions of water around sidechains that form interactions with other sidechains. (A-C) RDFs centered on the Tyr 6-carbon ring that forms cation-π, π-π, and amino-π interactions. (D) RDFs centered on Asp oxygens that form salt bridges. Insets show the approach of radial distribution functions to 1.
